# Ciliary neurotrophic factor curbs palmitate-induced inflammatory and metabolic changes in NPY/AgRP hypothalamic neurons

**DOI:** 10.64898/2026.10.02.756180

**Authors:** Jessica Perugini, Edoardo Scopini, Chiara Galli, Martina Senzacqua, Grandin Gaia, Georgia Colleluori, Ilenia Severi, Antonio Giordano

## Abstract

**Introduction:** In both humans and experimental animals, obesity is associated with hypothalamic inflammation and neuronal metabolic dysfunction, which contribute to disrupt body energy homeostasis. Although ciliary neurotrophic factor (CNTF) is a neuroprotective cytokine with recognized anti-obesity effects in peripheral metabolic tissues, its role in obesity-associated hypothalamic dysfunction is still poorly characterized. We investigated whether CNTF exerts any direct effects on mHypoE-N41 (N41) hypothalamic neurons and whether it modulates the effects induced by palmitate (PA), an established model of obesity-associated neuronal metabolic dysfunction.

**Methods:** N41 cells were administered CNTF alone or combined with PA to assess whether CNTF affects PA-induced lipotoxic stress. Gene and protein expression was assessed by qRT-PCR, western blotting, and immunofluorescence analysis.

**Results:** In N41 cells, CNTF activated the STAT3, ERK and AMPK signaling pathways. It also upregulated genes involved in mitochondrial metabolism and fatty acid oxidation, including *Ucp2*, *Cpt1α* and *Pgc1α*, as well as the proliferation-associated markers Ki-67 and PCNA. PA treatment induced inflammatory and endoplasmic reticulum stress responses and impaired insulin signaling. In PA-treated N41 cells, CNTF significantly attenuated the expression of pro-inflammatory markers *Tnfα, Mcp1* and *Nfkb*, it restored insulin-induced AKT phosphorylation, and partially reversed PA-induced changes in the expression of metabolic genes *Ucp2* and *Cpt1α*.

**Conclusions:** Collectively, these findings indicate that CNTF exerts direct protective metabolic effects on hypothalamic N41 neurons subjected to lipotoxic stress and supports a role for it in the preservation of neuronal metabolic function under obesogenic conditions.

## 1. Introduction

Ciliary neurotrophic factor (CNTF) is a member of the interleukin-6 family that signals through a tripartite receptor complex consisting of CNTF receptor alpha (CNTFRα), leukemia inhibitory factor receptor beta, and glycoprotein 130^1,2^. CNTF is produced primarily by astrocytes and tanycytes in the central nervous system and by Schwann cells in the peripheral nervous system. Initially characterized for its neurotrophic properties^3–5^, CNTF has eventually been recognized as a pleiotropic factor that is released in response to tissue injury and cellular stress^6–9^.

The metabolic relevance of CNTF in humans first emerged from clinical studies, originally designed to assess its therapeutic potential in amyotrophic lateral sclerosis, which found that treatment with recombinant CNTF unexpectedly resulted in weight loss^10,11^. Studies of experimental animals showed that recombinant CNTF induces weight loss and improves obesity-associated hyperglycemia, hyperinsulinemia, and hyperlipidemia through direct metabolic effects. Such effects are exerted on peripheral organs^12,13^ and include stimulation of adipose tissue lipolysis^14,15^, activation of brown adipose tissue–dependent non-shivering thermogenesis^16^, and protection of human multipotent adipose-derived stem (hMADS) adipocytes from tumor necrosis factor α (TNFα)-induced inflammation and insulin resistance^17^. Moreover, exogenous CNTF enhances fatty acid oxidation, it reverses obesity-induced insulin resistance in skeletal muscle^18,19^ and reduces high fat-induced hepatic steatosis while improving liver insulin responsiveness^20^. Importantly, individuals with obesity displayed higher circulating CNTF levels than healthy controls, suggesting that also in humans CNTF may be released in response to tissue stress, inflammation, and metabolic dysfunction, possibly acting as a protective factor against obesity-related metabolic complications^21,22^. Consistent with these findings, circulating CNTF declines in obese subjects following weight loss interventions^21^.

The hypothalamus plays a pivotal role in regulating energy metabolism by sensing and responding to changes in circulating peripheral signals including hormones, such as leptin and insulin, and nutrients, such as glucose and fatty acids^23–25^. In rodents, diet-induced obesity induces inflammatory and cellular stress responses within the hypothalamus, leading to neuronal injury and functional alterations in hypothalamic circuits involved in the control of energy balance^26,27^. Importantly, neuroimaging and *post mortem* studies also found hypothalamic inflammation and gliosis in humans with obesity, particularly in the mediobasal hypothalamus, strengthening the translational relevance of findings from experimental models^26,28,29^. Overfeeding, particularly diets enriched in saturated fatty acids such as palmitate (PA), promotes ectopic lipid accumulation and leads to the activation of pro-inflammatory pathways in hypothalamic neurons and glial cells, thereby contributing to low- grade hypothalamic inflammation and gliosis^26,30,31^. Consequently, hypothalamic inflammation impairs insulin signaling in hypothalamic neurons and alters the expression of feeding-related neuropeptides, disrupting the central regulation of energy homeostasis^31,32^. Importantly, experimental evidence suggests that modulation of hypothalamic inflammatory pathways may curb the metabolic complications of diet-induced obesity^33,34^.

Administered CNTF activates the Janus kinase/signal transducer and activator of transcription 3 (JAK/STAT3) signaling pathway in both orexigenic neuropeptide Y/agouti-related peptide (NPY/AgRP) and anorexigenic pro-opiomelanocortin (POMC) neurons in the arcuate nucleus (ARC) of the hypothalamus, exerting leptin-like effects^35–37^. In this regard, we found that CNTF preferentially activates JAK/STAT3 signaling in NPY/AgRP neurons located in the ARC ventromedial portion, close to the median eminence, an area characterized by a less restrictive blood– brain barrier^38^. CNTF also stimulated the mitogen-activated protein kinase/extracellular signal- regulated kinase (MAP/ERK) signaling pathway in β1/β2 tanycytes, specialized ependymal cells lining the floor of the third ventricle and interfacing with NPY neurons^38^. Altogether, our data^38^ suggested that CNTF administration may promote leptin entry and signaling in hypothalamic feeding centers. However, CNTF is produced in the hypothalamus, mainly by ependymal cells and tanycytes^39^, where under high-fat diet conditions both CNTF and its receptor CNTFRα are upregulated, whereas calorie restriction induces the opposite effect^40^. These observations suggest that, in states of caloric overload, tanycyte-derived CNTF may be secreted, or oversecreted, and act locally in the ARC, particularly on NPY/AgRP neurons, which are highly sensitive to the cytokine.

We analyzed mHypoE-N41 (N41) hypothalamic neurons exposed to CNTF, alone or combined with PA, a well-established *in vitro* model of obesity-associated hypothalamic neuroinflammation and neuronal metabolic dysfunction^41–43^, to establish *i)* whether CNTF exerts any direct effects on hypothalamic neurons and *ii)* whether it is able to counteract PA-induced lipotoxicity.

## 2. Materials and methods

### 2.1 Cell culture and reagents

The mouse hypothalamic cell line N41(RRID:CVCL_D442), which is derived from mouse hypothalamic primary cultures immortalized by retroviral transfer of SV40 T–Ag, was purchased from Cedarlane Corporation (Burlington, Canada). This clonal cell line has been assessed for the expression of a wide, and still-expanding, array of neuropeptides, enzymatic markers and biologically active receptors, and is a highly valuable tool to study cellular mechanisms and metabolic pathways involved in hypothalamic function. Information regarding the markers expressed in this cell line can be found on the Cedarlane website (https://www.cedarlanelabs.com). The cells were cultured in Dulbecco’s modified Eagle’s medium (DMEM; Euroclone, Milano, Italy), supplemented with 10% fetal bovine serum (Thermo Fisher Scientific, Waltham, MA, USA) and 1% penicillin-streptomycin (Euroclone), and maintained at 37 °C in a humidified atmosphere containing 5% CO_2_. Mouse recombinant CNTF was purchased from PeproTech (London, UK), whereas PA and insulin were obtained from Sigma-Aldrich (Milano, Italy). PA was prepared according to the protocol described by Belsham et al.^41^. Experiments were performed on N41 cells grown to 80-90% confluence. Treatment duration and CNTF, PA and insulin concentrations are reported in the Results section.

### 2.2 qRT-PCR

Total RNA was extracted from N41 cells using TRIzol reagent (Invitrogen, Carlsbad, CA, USA), purified, digested with ribonuclease-free deoxyribonuclease, and concentrated using RNeasy Micro kit (Qiagen, Milano, Italy) according to the manufacturer’s instructions. For the determination of mRNA levels, 1 μg of total RNA was reverse-transcribed using the High-Capacity cDNA Reverse Transcription Kit with RNase Inhibitor (Applied BioSystems, Foster City, CA, USA) in a total volume of 20 μL. Quantitative real-time PCR (qRT-PCR) was performed using gene-specific primers (Table 1A) and PowerUp™ SYBR™ Green Master Mix (Applied BioSystems), according to the manufacturer’s instructions. Reactions were carried out in a StepOnePlus™ Real-Time PCR System (Applied BioSystems) using 50 ng of cDNA in a final reaction volume of 10 μL. A control reaction without reverse transcriptase in the amplification mixture was run for each sample to rule out genomic DNA contamination.

**Table 1A.**
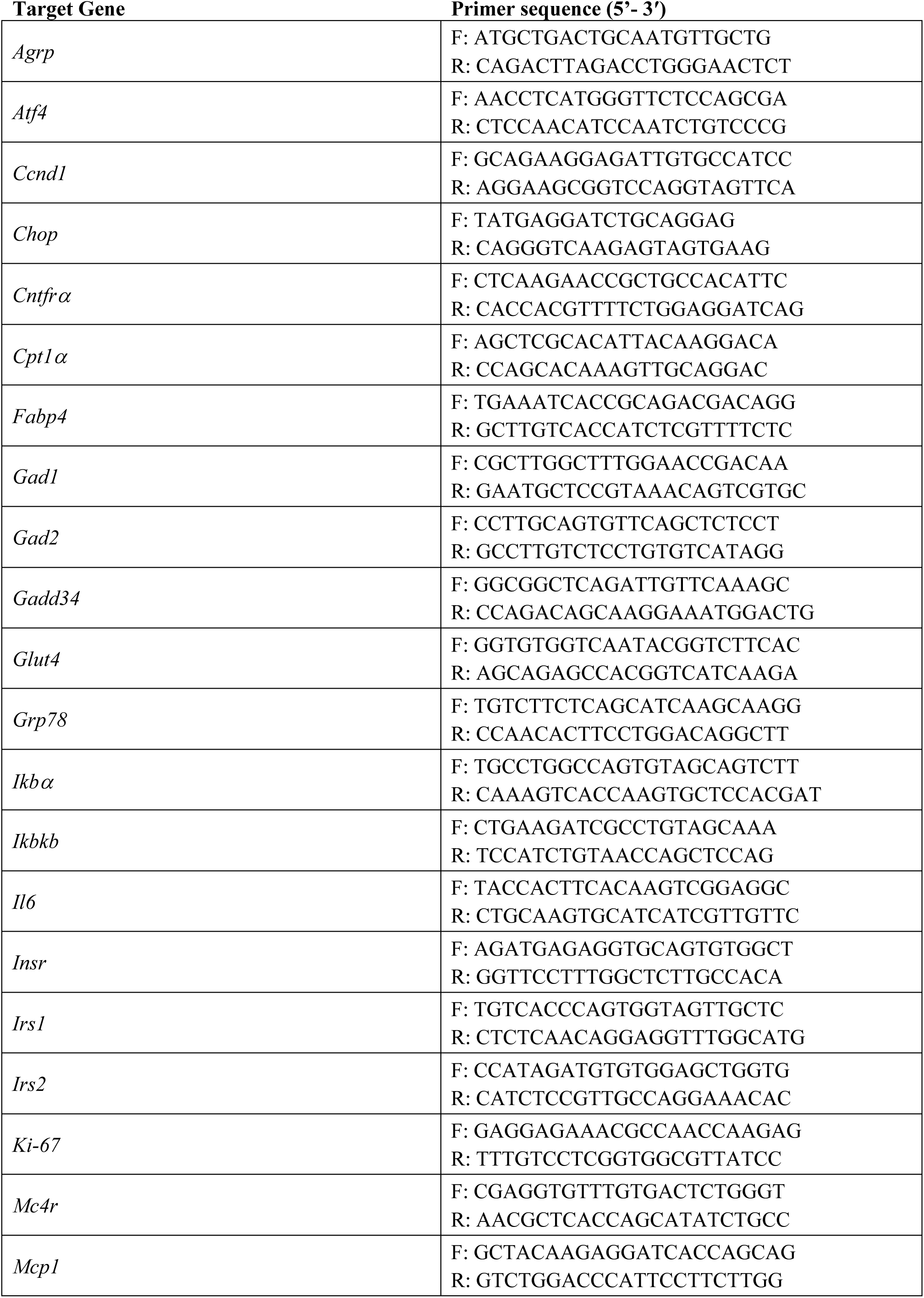
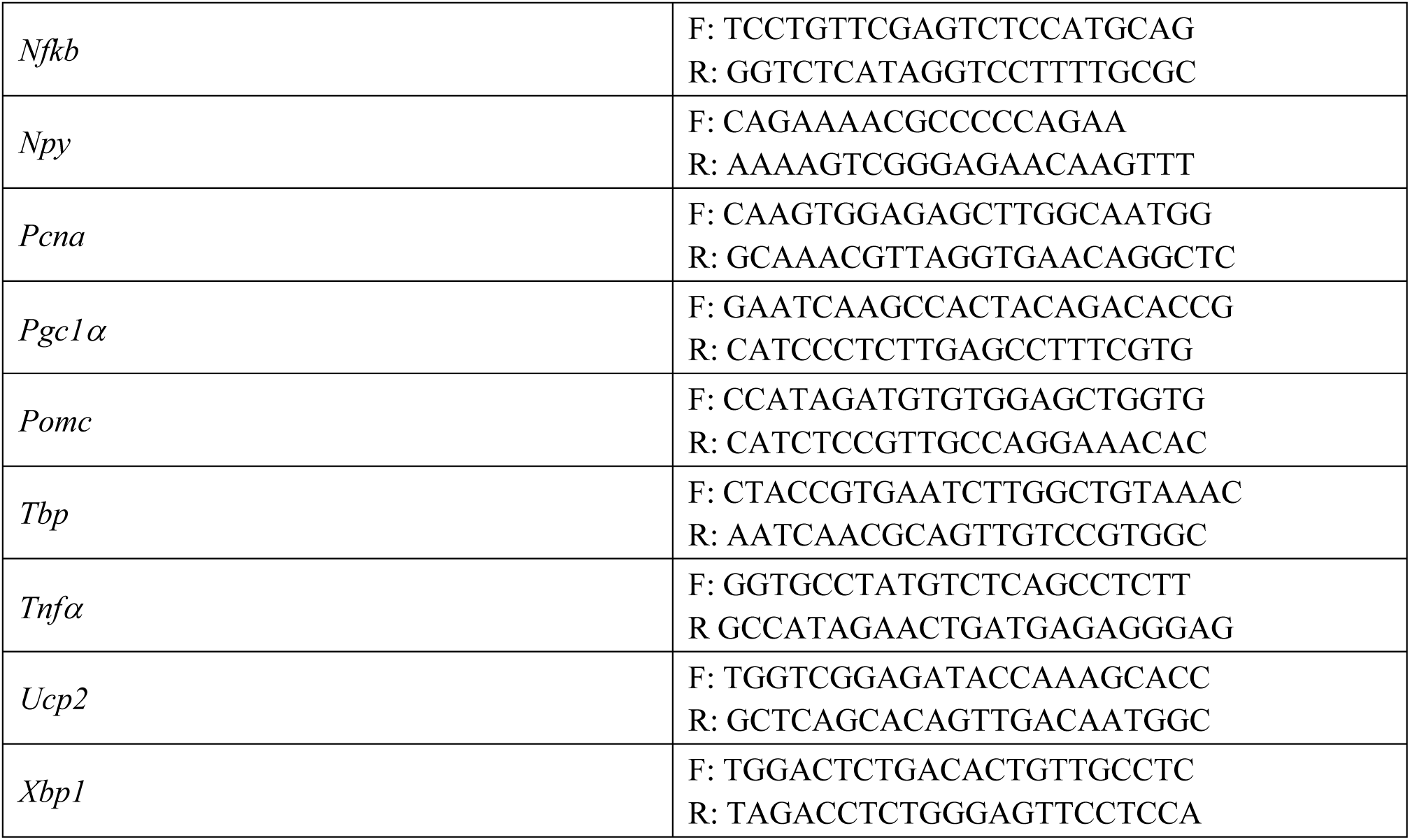
Primer list for qRT-PCR (all from Invitrogen #10629186).

No-template controls were included in all experiments as negative controls. TATA box-binding protein (*Tbp*) was the endogenous control for normalization of gene expression. Relative mRNA expression was determined by the ΔCt method (2^-ΔCt^). Where applicable, data are presented as fold change relative to untreated cells calculated by the ΔΔCt method (2^-ΔΔCt^).

### 2.3 Immunofluorescence and confocal microscopy

For immunofluorescence experiments, N41 cells were cultured in two-well Lab-Tek Chamber Slides (Nalge Nunc International, Naperville, IL, USA), washed with phosphate-buffered saline (PBS, pH 7.4), fixed with 4% paraformaldehyde in PBS for 10 min at room temperature (RT), and permeabilized with PBS containing 0.1% Triton X-100 for 10 min. After three washes in PBS, cells were blocked with 10% normal donkey serum (Jackson Immuno Research, West Grove, PA, USA) in PBS, then incubated overnight at 4 °C with a mixture of two primary antibodies (Table 1B). The next day, cells were thoroughly rinsed with PBS and incubated for 1 h at RT with a cocktail of fluorophore- conjugated secondary antibodies (Table 1C). After washing with PBS, nuclear staining was performed using TO-PRO™-3 Iodide (Invitrogen). Finally, cells were air-dried and coverslipped with Vectashield mounting medium (Vector Laboratories, Burlingame, CA, USA). Images were acquired using a motorized Leica DM6000 microscope at different magnifications. Fluorescence was detected with a Leica TCS-SL spectral confocal microscope (Wetzlar, Germany) equipped with an Argon and He/Ne mixed gas laser. Fluorophores were excited at 488, 543, and 649 nm and imaged separately. Images (1,024 × 1,024 pixels) were obtained sequentially from three channels using a confocal pinhole of 1.1200 and stored as TIFF files. When the primary antibody was omitted, staining was never observed. For morphometric analysis, 10 non-overlapping fields were randomly selected for each experimental condition and imaged at 40x magnification. The total number of nuclei, identified by TO-PRO-3 staining, and the number of Ki67-positive nuclei were counted in each field using ImageJ/Fiji software (RRID:SCR_003070), and the percentage of Ki-67-positive nuclei was determined. Values calculated in the 10 fields were averaged to generate a single value for each experimental condition

**Table 1B.**
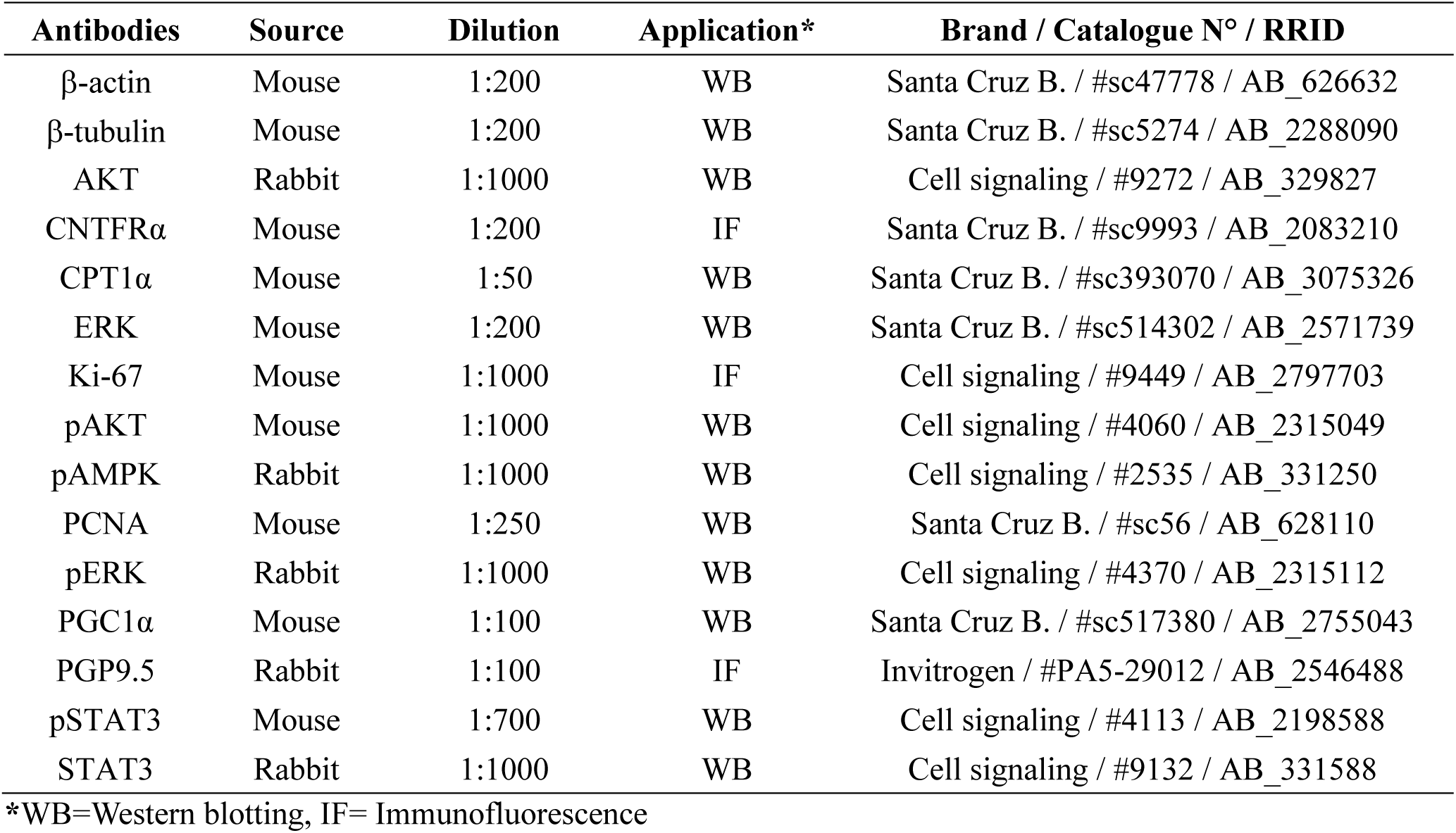
Primary antibodies.

**Table 1C.**
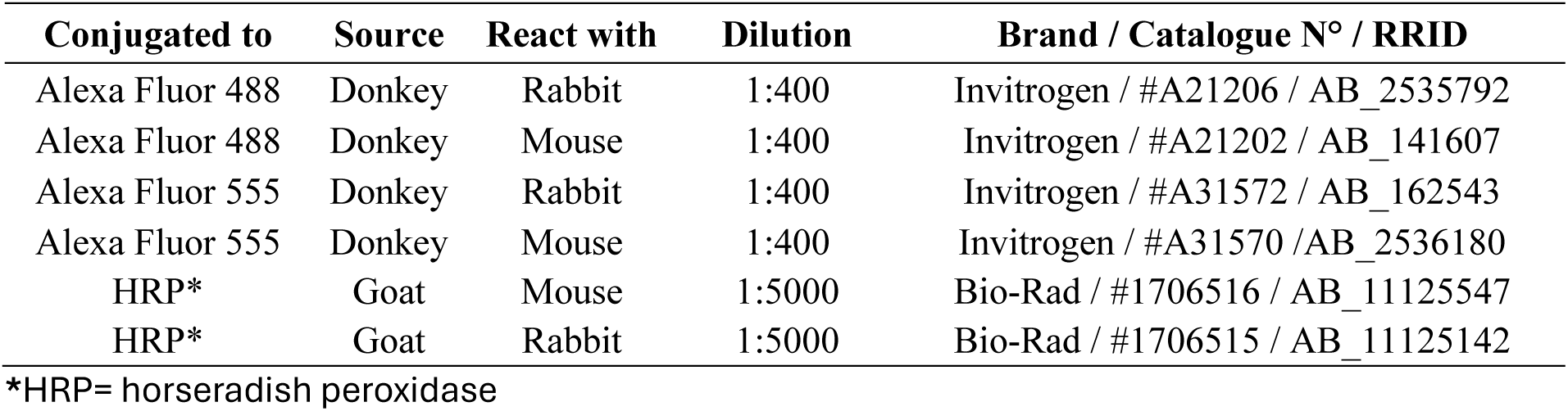
Secondary antibodies.

### 2.4 Western blotting

For western blotting experiments, cell lysates were prepared using a lysis buffer containing 50 mM Tris–HCl (pH 7.4), 1% NP-40, 1 mM EDTA, 150 mM NaCl, 1 mM sodium orthovanadate, 0.5% sodium deoxycholate, 0.1% SDS, 1 mM phenylmethylsulfonylfluoride, and 2 µg/mL aprotinin in deionized water. Samples were centrifuged and protein concentrations determined by the Bradford Protein Assay (Bio-Rad Laboratories, Segrate, Italy). Equal amounts of protein were separated by SDS-PAGE and transferred onto nitrocellulose membranes using the Trans-Blot Turbo^TM^ Transfer system (Bio-Rad). To check loading and transfer efficiency, membranes were visualized with Ponceau S solution (Santa Cruz Biotechnology, Santa Cruz, CA, USA). Membranes were then blocked for 1 h at RT in TBS-T (50 mM Tris-HCL [pH 7.6], 200 mM NaCl, and 0.1% Tween-20) containing 5% non-fat dried milk, followed by overnight incubation at 4 °C with the primary antibody (Table 1B). After washing in TBS-T, membranes were incubated for 1 h at RT with the appropriate horseradish peroxidase-conjugated secondary antibody (Table 1C). Immunoreactive bands were visualized using the Clarity™ Western ECL substrate and the Chemidoc Imaging System (all from Bio-Rad). Band densitometric analysis was performed using Bio-Rad Image Lab software (RRID:SCR_014210). Where applicable, membranes were stripped, washed, and re-probed for total protein content.

### 2.5 Statistical analysis

Data are presented as mean ± standard error (SEM) and were analyzed by GraphPad Prism software (RRID:SCR_002798). All experiments were run in triplicate. Statistical significance was assessed by one-way analysis of variance (ANOVA) followed by Tukey’s multiple comparisons *post hoc* test. Differences were considered statistically significant at *p* < 0.05.

## 3. Results

### 3.1 Characterization of N41 cells

By light microscopy, N41 cells displayed a neuron-like morphology with small cell bodies and multiple, elongated cytoplasmic processes (Figure 1A). Gene expression analysis confirmed the expression of *Npy* and *Agrp*. Cells also expressed low levels of glutamate decarboxylase 1 (*Gad1*), one of the enzymes responsible for GABA synthesis, consistent with the GABAergic phenotype of NPY/AgRP neurons (Figure 1B). Although multiple GABAergic neuronal populations in hypothalamic circuits contribute to the regulation of feeding behavior, NPY/AgRP neurons of the ARC are a key orexigenic population whose function is to integrate peripheral metabolic signals, playing a crucial role in driving hunger and food intake^44,45^. In contrast, ARC POMC neurons are a functionally opposing anorexigenic population, a substantial subset of which display a glutamatergic phenotype^46^. Additionally, melanocortin-4 receptor (*Mc4r*) was barely detectable whereas the expression of *Pomc* and glutamate decarboxylase 2 (*Gad2*) was undetectable (Figure 1B). Overall, the expression of *Npy*, *Agrp*, and *Gad1*, coupled with the absence of *Pomc*, supports the use of N41 cells as an *in vitro* model predominantly reflecting the molecular phenotype of NPY/AgRP hypothalamic neurons. N41 cells also expressed several key components of the insulin signaling pathway, including insulin receptor (*Insr*), insulin receptor substrates 1 and 2 (*Irs1* and *Irs2*), and glucose transporter 4 (*Glut4*), as well as markers of fatty acid signaling and β-oxidation, such as fatty acid-binding protein 4 (*Fabp4*), carnitine palmitoyltransferase 1α (*Cpt1α*), and uncoupling protein 2 (*Ucp2*) (Figure 1B).

**Figure 1.**
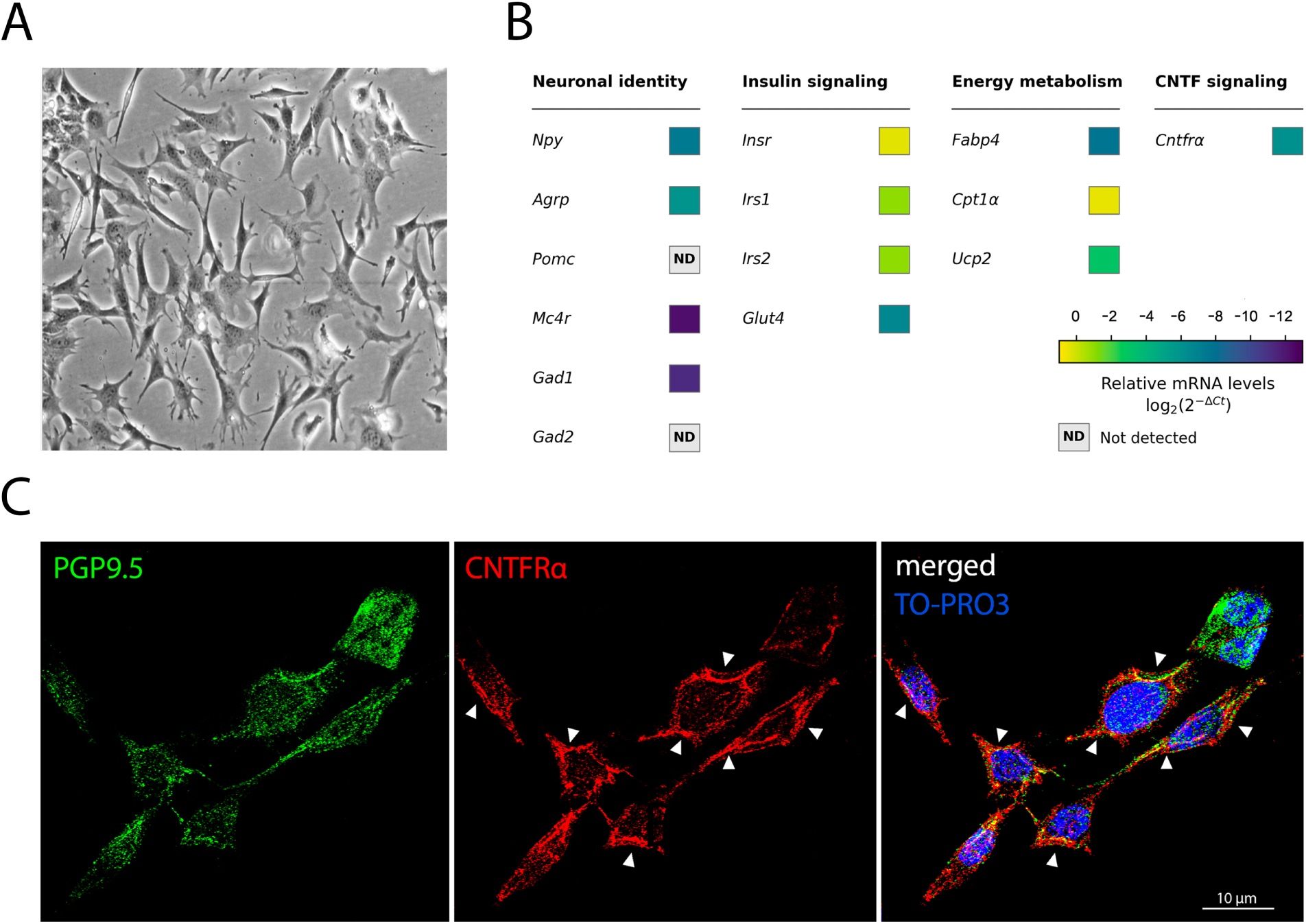
Characterization of N41 cells. (A) Representative light microscopy image showing cell morphology. (B) Basal gene expression of neuronal (*Npy*, *Agrp, Pomc, Mc4r*, *Gad1* and *Gad2*), insulin signaling (*Insr*, *Irs1*, *Irs2* and *Glut4*), and energy metabolism-related (*Fabp4*, *Cpt1a* and *Ucp2*) genes and of *Cntfrα* in N41 cells. Gene expression was determined by RT-qPCR and expressed as relative mRNA levels (2^-ΔCt^) normalized to *Tbp*. Heatmap color intensity represents log_₂_- transformed relative mRNA expression. ND, not detected. (C) Representative double immunostaining confocal microscopy images showing PGP9.5 staining (left, green) and CNTFRα staining (middle, red). TO-PRO3 nuclear counterstaining is shown in blue. CNTFRα immunoreactivity was detected throughout the cytoplasm and along the plasma membrane (white arrowheads).

To assess the responsiveness of hypothalamic N41 neurons to CNTF, we first tested the expression of its specific receptor, CNTFRα. Gene expression analysis revealed detectable levels of *Cntfrα* mRNA (Figure 1B). Double immunofluorescence confocal microscopy further confirmed the expression of CNTFRα protein in PGP9.5-positive N41 neurons. CNTFRα immunoreactivity was detected throughout the cell body but especially on the plasma membrane, consistent with the localization of receptors for secreted molecules in mammalian cells (Figure 1C).

Taken together, their expression profile indicates that N41 cells are a suitable *in vitro* model of NPY/AgRP-expressing hypothalamic neurons to investigate metabolic regulation and CNTF-mediated effects.

### 3.2 In N41 neurons, CNTF activates its canonical intracellular signaling pathways

To assess whether N41 neurons are responsive to CNTF, we evaluated the activation of its canonical downstream signaling pathways, JAK2/STAT3 and MAPK/ERK^2,47^. To do this, cells were treated with increasing concentrations of recombinant CNTF protein (0.1, 1, and 10 nM) for 15 min. Western blot analysis disclosed dose-dependent phosphorylation of both STAT3 and ERK (Figure 2A). Densitometric analysis revealed no significant activation with 0.1 nM CNTF, whereas treatment with 1 and 10 nM CNTF induced robust phosphorylation of both signaling proteins, confirming activation of the JAK2/STAT3 (Figure 2B) and MAPK/ERK pathways (Figure 2C). To gain further insight into the temporal dynamics of CNTF signaling, cells were exposed to 1 nM CNTF for 15, 30, or 60 min before western blot analysis (Figure 2D). Phosphorylation of STAT3 (Figure 2E) and ERK (Figure 2F) showed a significant peak at 15 min followed by a gradual decline, reflecting the transient activation of the two pathways. These findings demonstrate that CNTF activates well-established CNTF-responsive signaling pathways^2^ in N41 neurons, supporting their use as an *in vitro* model to investigate the cellular effects of CNTF on orexigenic hypothalamic neurons.

**Figure 2.**
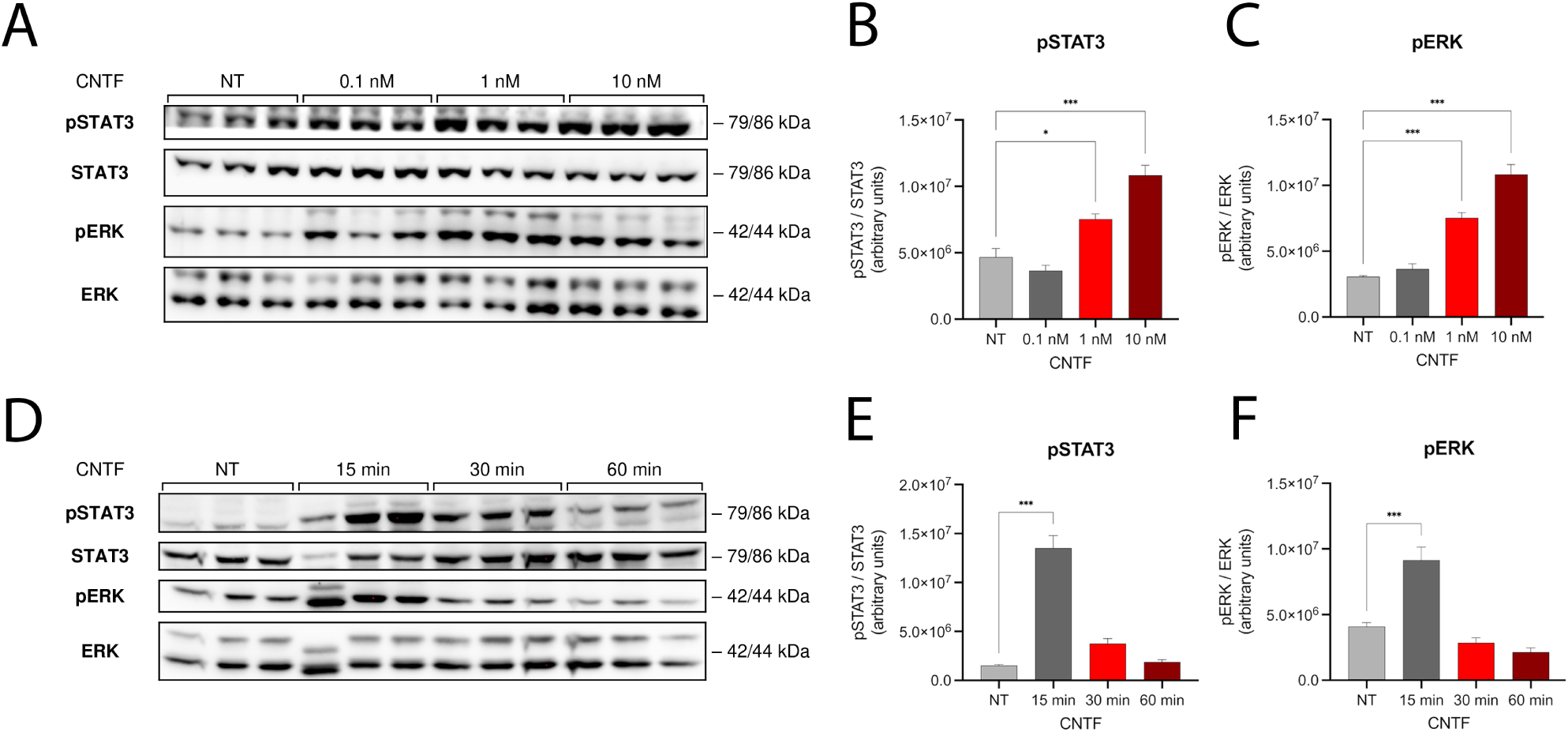
JAK/STAT3 and MAPK/ERK signaling pathways in N41 hypothalamic neurons after CNTF treatment. (A) Representative immunoblots and densitometric quantification of the dose-dependent STAT3 (B) and ERK (C) phosphorylation in N41 neurons treated with increasing concentrations of CNTF for 15 min. Representative immunoblots (D) and densitometric quantification of time-dependent STAT3 (E) and ERK (F) phosphorylation in N41 neurons treated with 1 nM CNTF for the indicated times. Densitometric values were normalized to total STAT3 and ERK levels, respectively. Data are mean ± SEM from three 3 independent experiments. *p < 0.05; **p < 0.01; ***p < 0.001 vs. untreated cells (NT), as determined by one-way ANOVA followed by Tukey’s post hoc test.

### 3.3 In N41 neurons, CNTF modulates proliferation-associated markers and neuropeptide expression

In addition to its metabolic actions^17–19^, CNTF has been suggested to promote adult hypothalamic neurogenesis, a process involved in energy homeostasis^48–50^. Since constitutive neural cell proliferation persists in the adult hypothalamus^48^, we investigated whether CNTF modulates the expression of proliferation-associated markers in N41 hypothalamic neurons. Gene expression analysis revealed significant *Ki-67* upregulation following CNTF treatment at both 6 and 24 h compared with untreated cells, whereas proliferating cell nuclear antigen (*Pcna*) and cyclin D1 (*Ccnd1*) expression levels were not significantly affected (Figure 3A). We further assessed the effects of CNTF on proliferation at the protein level by analyzing PCNA expression by western blotting. The blots showed a significant increase in PCNA protein levels 6 h into CNTF treatment (Figure 3B and Figure 3C). Consistent with these findings, immunofluorescence analysis demonstrated an increased number of Ki-67–positive nuclei at the same time point (Figure 3D). Quantitative morphometric analysis demonstrated a significant increase in the percentage of Ki-67–positive cells compared with untreated cells (Figure 3E), suggesting activation of a proliferation-related program.

**Figure 3.**
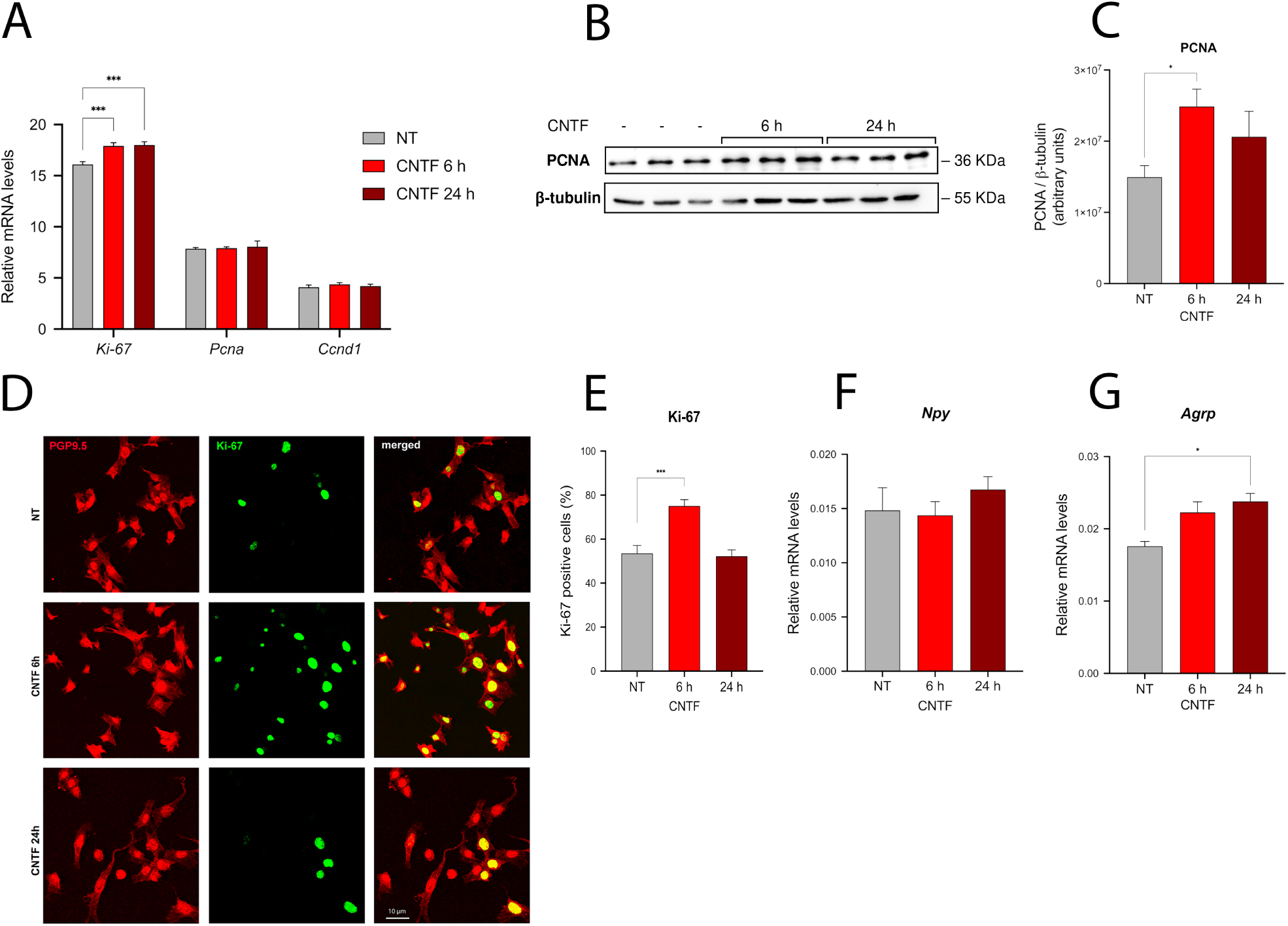
Proliferation-associated and orexigenic neuronal markers in N41 hypothalamic neurons after CNTF treatment. (A) Relative mRNA expression of *Ki-67*, *Pcna* and *Ccnd1* in N41 neurons treated with 1 nM CNTF for 6 or 24 h. Gene expression was determined by RT-qPCR and expressed as relative mRNA levels (2^-ΔCt^) normalized to *Tbp*. (B) Representative immunoblot and densitometric quantification of PCNA protein levels (C) in N41 neurons treated with 1 nM CNTF for 6 or 24 h. PCNA protein levels were normalized to β-tubulin. (D) Representative double immunostaining confocal microscopy images showing PGP9.5 staining (red) and Ki-67 staining (green) in the nuclei of untreated (NT) and CNTF-treated N41 neurons (1 nM, 6 or 24 h). Merged images in right panels. Scale bar, 10 μm. (E) Quantification of Ki-67-positive cells (%). Relative mRNA expression of the orexigenic neuropeptides *Npy* (F) and *Agrp* (G) in N41 neurons treated with 1 nM CNTF for 6 or 24 h. Gene expression was determined by RT-qPCR and expressed as relative mRNA levels (2^-ΔCt^) normalized to *Tbp*. Data are mean ± SEM from three independent experiments. *p < 0.05; **p < 0.01; ***p < 0.001 vs. untreated cells (NT), as determined by one-way ANOVA followed by Tukey’s post hoc test.

To achieve further characterization of the effects of CNTF on the N41 neuronal phenotype, we assessed the expression of neuropeptides associated with hypothalamic neuron differentiation. Gene expression analysis disclosed that *Npy* levels were not significantly affected by CNTF at any time point (Figure 3F), whereas *Agrp* was significantly upregulated after 24 h of CNTF exposure (Figure 3G). These findings suggest that, besides affecting proliferation-associated markers, CNTF may selectively modulate the expression of neuropeptides linked to the orexigenic neuronal phenotype.

### 3.4 In N41 neurons, CNTF attenuates PA-induced inflammation

To establish an *in vitro* model of neuroinflammation in N41 neurons, we assessed their response to PA, a long-chain saturated fatty acid. To do this, we exposed cells to 200 μM PA, a concentration that does not affect viability^41^, for 3, 6, or 24 h. At 24 h, PA induced a significant increase in the expression of pro-inflammatory genes, including *Il6*, *Tnfα*, *Mcp1*, *Nfkb*, and *Ikbkb*, indicating the onset of an inflammatory response in N41 neurons (Figure 4A). Notably, *Mcp1*, a chemokine involved in monocyte recruitment, was already significantly upregulated at 6 h. At 24 h, PA treatment also caused a marked increase in *Chop*, a key marker of endoplasmic reticulum (ER) stress involved in apoptotic signaling^51^. Consistent with the latter finding, PA induced significant upregulation of other ER stress– related genes^51^, namely *Gadd34*, *Xbp1* and *Atf4* at 6 and 24 h, and *Grp78* at 24 h (Figure 4A). Collectively, these findings indicate that PA exposure elicits inflammatory and stress-related responses in N41 neurons.

**Figure 4.**
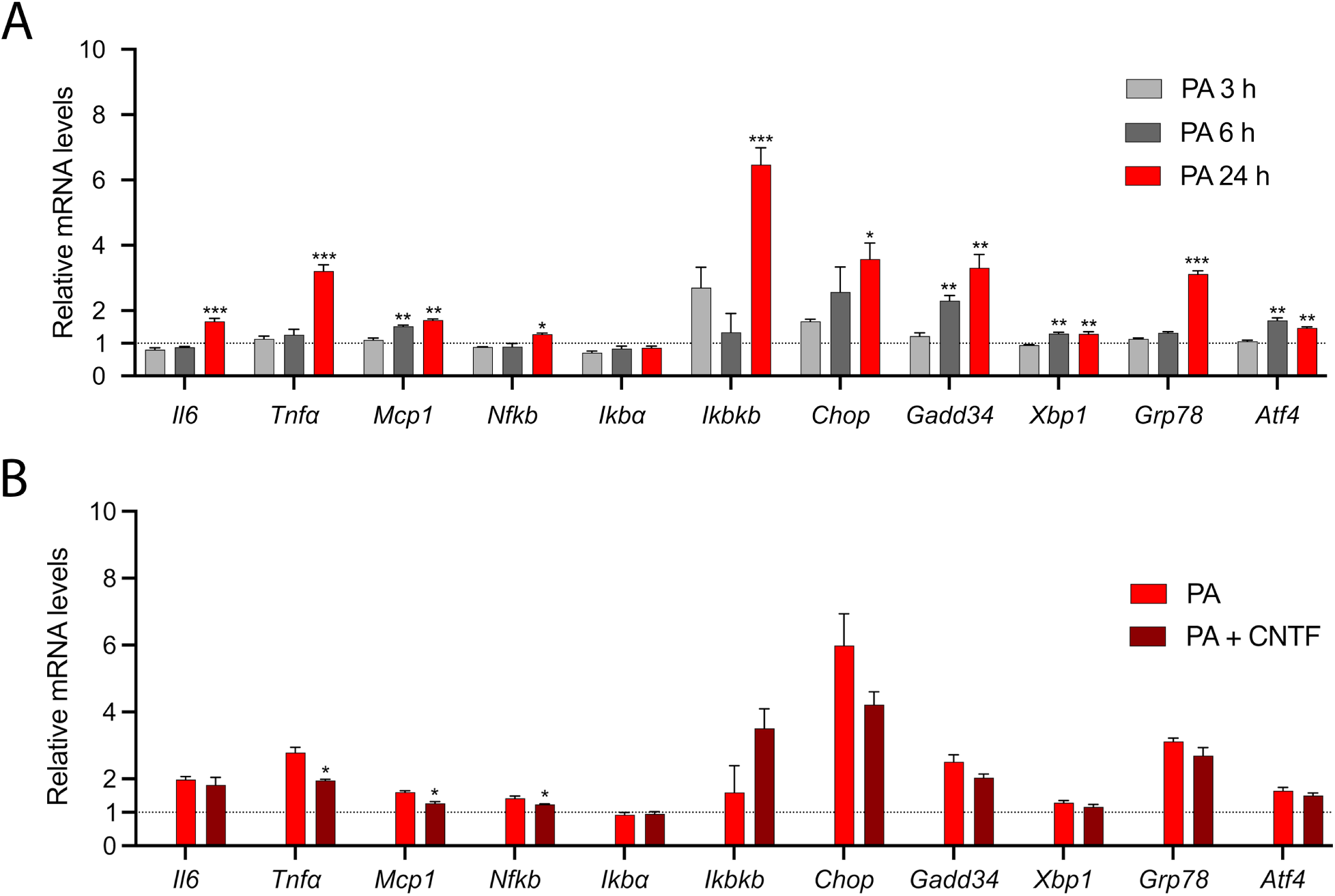
Inflammatory and endoplasmic reticulum stress-related genes in N41 hypothalamic neurons after PA and CNTF treatment. (A) Relative mRNA expression of inflammatory (*Il6*, *Tnfa*, *Mcp1*, *Nfkb*, *Ikba* and *Ikbkb*) and ER stress-related (*Chop*, *Gadd34*, *Xbp1*, *Grp78* and *Atf4*) genes in N41 neurons treated with 200 μM PA for 3, 6 or 24 h. *p < 0.05; **p < 0.01; ***p < 0.001 vs. untreated (NT) cells, as determined by one-way ANOVA followed by Tukey’s post hoc test. (B) Relative mRNA expression of inflammatory and ER stress-related genes in N41 neurons co-treated with 200 μM PA and 1 nM CNTF for 24 h. Gene expression was determined by RT-qPCR and expressed as fold change relative to untreated (NT) cells using the 2^-ΔΔCt^ method, with *Tbp* as the reference gene. Data are mean ± SEM from three independent experiments. *p < 0.05; **p < 0.01; ***p < 0.001 vs. PA treated (PA) cells, as determined by one-way ANOVA followed by Tukey’s post hoc test.

Based on its known anti-inflammatory effects^17^, we next evaluated the ability of CNTF to *i)* offset PA-induced upregulation of pro-inflammatory genes and *ii)* modulate PA-induced ER stress in N41 neurons co-treated with PA for 24 h. We found that CNTF significantly curbed the PA-induced upregulation of *Tnfα*, *Mcp1*, and *Nfkb* mRNA levels (Figure 4B). In addition, CNTF reduced the expression of ER stress–related genes, including *Chop*, *Gadd34*, and *Grp78*, although the reduction did not reach statistical significance (Figure 4B). These findings suggest that CNTF may attenuate some inflammatory responses triggered by PA, lending support to a protective role for it under lipotoxic conditions.

### 3.5 In N41 neurons, CNTF enhances insulin-induced AKT phosphorylation and attenuates PA-induced insulin resistance

To investigate the effects of CNTF on the insulin signaling pathway, we first examined whether it modulates AKT phosphorylation in N41 neurons under basal conditions. Based on western blot analysis, neither 1 nor 10 nM CNTF (20 min) significantly affected AKT phosphorylation at Ser473 (Figure 5A and Figure 5C), indicating that CNTF does not activate the AKT pathway under basal conditions. We next evaluated whether CNTF modulates insulin responsiveness by treating N41 neurons with CNTF (1 or 10nM, 20 min) and insulin (100 nM, 20 min). Co-treatment significantly enhanced insulin-induced AKT phosphorylation compared with insulin treatment alone (Figure 5B and Figure 5C). Given the central role of GLUT4 in insulin-responsive neuronal glucose uptake and hypothalamic glucose sensing, we next examined whether CNTF modulated *Glut4* expression^52^ and found that CNTF significantly increased *Glut4* mRNA expression at 24 h (Figure 5D). Acute insulin stimulation (100 nM, 10 min) prior to cell lysis did not significantly affect *Glut4* gene expression, in line with reports of its rapid action on GLUT4 trafficking rather than on transcriptional regulation ^53^. These findings indicate that CNTF enhances AKT signaling and increases the expression of a key component of neuronal glucose metabolism.

**Figure 5.**
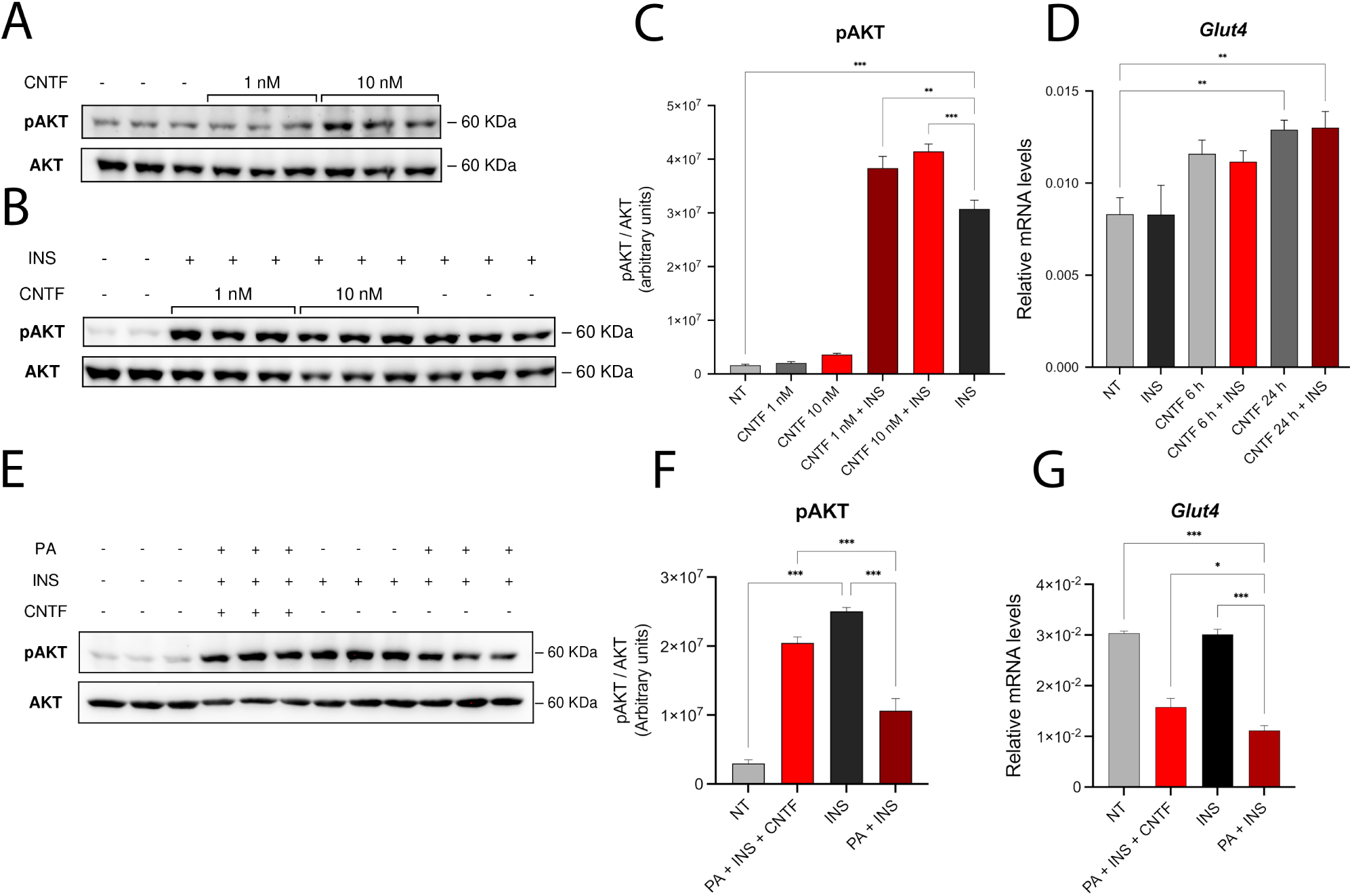
AKT signaling in N41 hypothalamic neurons after CNTF treatment. (A) Representative immunoblots showing AKT phosphorylation in N41 neurons treated with CNTF (1 or 10 nM) for 20 min. (B) Representative immunoblots showing insulin-stimulated AKT phosphorylation in N41 neurons co-treated with CNTF (1 or 10 nM) and insulin (100 nM) for 20 min. (C) Densitometric quantification of 473-serine AKT phosphorylation. Phosphorylated AKT levels were normalized to total AKT. (D) Relative *Glut4* mRNA expression in N41 neurons treated with CNTF (1 nM) for 6 or 24 h, in the presence/absence of insulin (100 nM, 10 min). Gene expression was determined by RT-qPCR and expressed as relative mRNA levels (2^-ΔCt^) normalized to *Tbp*. (E) Representative immunoblot and (F) densitometric quantification of insulin-stimulated 473-serine AKT phosphorylation in N41 neurons co-treated with 200 μM PA and 1nM CNTF for 24 h, followed by insulin stimulation (100 nM, 10 min) where indicated. Phosphorylated AKT levels were normalized to total AKT. (G) Relative *Glut4* mRNA expression in N41 neurons co-treated with 200 μM PA and 1nM CNTF for 24 h, followed by insulin stimulation (100 nM, 10 min) where indicated. Gene expression was determined by RT-qPCR and expressed as relative mRNA levels (2^-ΔCt^) normalized to *Tbp*. Data are mean ± SEM from three independent experiments. *p < 0.05; **p < 0.01; ***p < 0.001 as determined by one-way ANOVA followed by Tukey’s post hoc test.

Several studies have found that PA induces insulin resistance in pancreatic β-cells, hepatocytes, adipocytes, and hypothalamic neurons^41,54–56^. Accordingly, we tested its ability to induce insulin resistance in N41 neurons. As expected, prolonged (24 h) exposure to PA and insulin stimulation (100 nM, 10 min) prior to cell lysis resulted in a significant reduction in insulin-induced AKT phosphorylation compared with insulin exposure alone (Figure 5E and Figure 5F), suggesting the development of insulin resistance. To determine whether CNTF was able to restore insulin sensitivity in PA-exposed neurons, N41 cells were co-treated with 200 μM PA and 1 nM CNTF for 24 h, followed by stimulation with insulin (100 nM, 10 min) prior to cell lysis. Notably, CNTF was able to counteract the reduction in insulin-induced AKT phosphorylation in PA-treated neurons, reflecting improved insulin signaling (Figure 5E and Figure 5F). Consistent with these findings, PA and insulin treatment reduced *Glut4* mRNA levels, whereas CNTF partially but significantly restored *Glut4* expression in PA-treated neurons (Figure 5G). Altogether, these data show that CNTF curbs PA-induced insulin resistance by enhancing AKT activation and *Glut4* mRNA levels, suggesting a protective role for CNTF in preserving insulin responsiveness in hypothalamic neurons exposed to lipotoxic conditions.

### 3.6 In N41 neurons, CNTF activates AMPK signaling and modulates the expression of genes associated with mitochondrial oxidative metabolism

AMP-activated protein kinase (AMPK) is a key hypothalamic energy sensor that regulates mitochondrial energy metabolism by promoting fatty acid oxidation, mitochondrial biogenesis, and oxidative metabolism^57^. Since CNTF activates AMPK in metabolically active peripheral tissues, such as skeletal muscle and adipose tissue^17,19^, we investigated whether AMPK signaling is also activated by CNTF in hypothalamic neurons. Western blot analysis revealed a significant increase in AMPK phosphorylation (pAMPK) 30 min and 60 min into CNTF treatment, indicating the activation of this energy-sensing pathway in N41 neurons (Figure 6A and Figure 6B).

**Figure 6.**
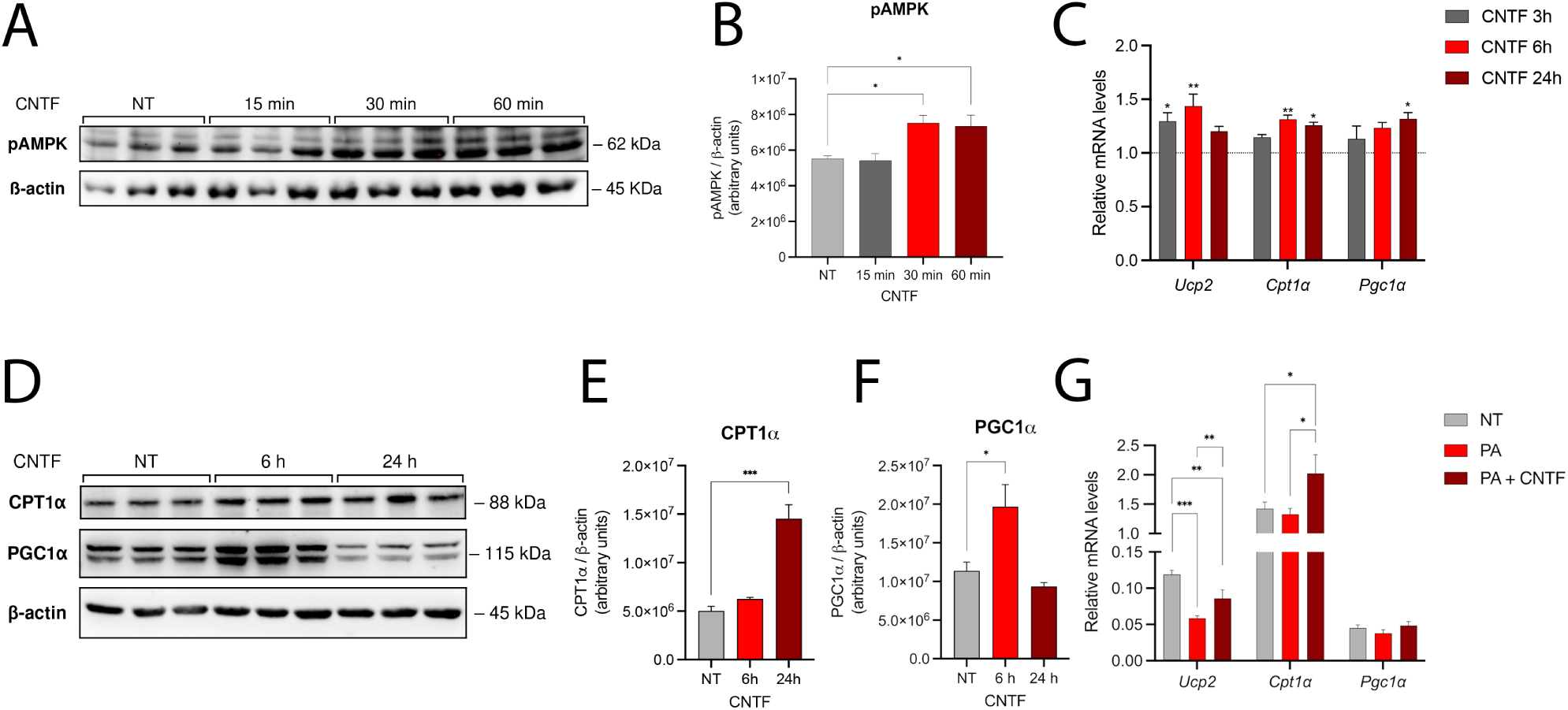
Energy metabolism-related markers in N41 hypothalamic neurons after CNTF and PA treatment. (A) Representative immunoblots and densitometric quantification of AMPK phosphorylation (B) in N41 neurons treated with CNTF (1 nM) for 15, 30 or 60 min. Phosphorylated AMPK levels were normalized to β-actin. (C) Relative mRNA expression of *Ucp2*, *Cpt1α* and *Pgc1α* in N41 neurons treated with CNTF (1 nM) for 3, 6 or 24 h. Gene expression was determined by RT- qPCR and expressed as fold change relative to untreated (NT) cells using the 2^-ΔΔCt^ method, with *Tbp* as the reference gene. *p < 0.05; **p < 0.01; ***p < 0.001 vs. untreated cells (NT), as determined by one-way ANOVA followed by Tukey’s post hoc test. (D) Representative immunoblots and densitometric quantification of CPT1α (E) and PGC1α (F) protein levels in N41 neurons treated with CNTF (1 nM) for 6 or 24 h. Protein levels were normalized to β-actin. (G) Relative mRNA expression of *Ucp2*, *Cpt1α* and *Pgc1α* in N41 neurons co-treated with 200 μM PA and 1nM CNTF for 24 h. Gene expression was determined by RT-qPCR and expressed as relative mRNA levels (2^-ΔCt^) normalized to *Tbp*. Data are mean ± SEM from three independent experiments. *p < 0.05; **p < 0.01; ***p < 0.001, as determined by one-way ANOVA followed by Tukey’s post hoc test.

Because AMPK has been implicated in the regulation of mitochondrial oxidative metabolism^57^, we next investigated whether CNTF modulates the expression of genes involved in this metabolic program^17,19^ in N41 hypothalamic neurons. CNTF upregulated the expression of genes associated with mitochondrial oxidative metabolism, including *Ucp2* at 3 and 6 h and *Cpt1a* at 6 and 24 h, whereas peroxisome proliferator activator receptor γ coactivator-1α (*Pgc1a*) was significantly upregulated only at 24 h (Figure 6C). Importantly, western blot analysis of N41 cell protein extracts (Figure 6D) confirmed increased CPT1α (Figure 6E) and PGC1α (Figure 6F) protein expression 24 and 6 h into CNTF treatment, respectively. Altogether, these findings indicate that in N41 hypothalamic neurons CNTF activates AMPK signaling and modulates the expression of genes related to mitochondrial function and fatty acid oxidation.

We next sought to determine whether these adaptive responses are impaired under lipotoxic conditions. For this purpose, N41 neurons were treated with 200 μM PA and 1 nM CNTF for 24 h. PA treatment significantly reduced *Ucp2* expression (Figure 6G). Given the role of UCP2 in maintaining mitochondrial function and limiting oxidative stress^58^, this reduction underscores the detrimental effects of saturated fatty acids on neuron metabolic homeostasis. Interestingly, CNTF attenuated the PA-induced downregulation of *Ucp2* expression (Figure 6G), suggesting a protective action for it on mitochondrial function under lipotoxic conditions. Notably, *Cpt1α* expression was not significantly affected by PA exposure. In contrast, combined PA and CNTF treatment significantly upregulated *Cpt1α* (Figure 6G), highlighting the ability of CNTF to promote mitochondrial fatty acid oxidation in a lipotoxic environment. Together with the restoration of *Ucp2* expression, the CNTF-induced upregulation of *Cpt1α* supports a coordinated mitochondrial adaptive response directed at limiting lipid-induced metabolic stress.

### 3.7 In N41 neurons, CNTF fails to modulate proliferation-associated markers and neuropeptide expression under lipotoxic conditions

Given that obesity-associated hypothalamic inflammation impairs hypothalamic cell proliferation and neurogenesis through neuroinflammatory mechanisms associated with metabolic dysfunctions, *e.g.* excessive caloric intake, glucose intolerance, insulin resistance, and overweight^59^, we investigated the impact of CNTF on proliferation-associated markers under lipotoxic conditions. Gene expression analysis demonstrated that PA treatment significantly reduced *Ki-67* mRNA levels compared with no treatment (Figure 7A), suggesting that lipotoxicity impairs the expression of this proliferation-associated marker. Although CNTF upregulated *Ki-67* under basal conditions, it failed to restore its expression in PA-treated neurons (Figure 7A). Neither PA nor CNTF significantly affected *Pcna* and *Ccnd1* mRNA levels (Figure 7A).

**Figure 7.**
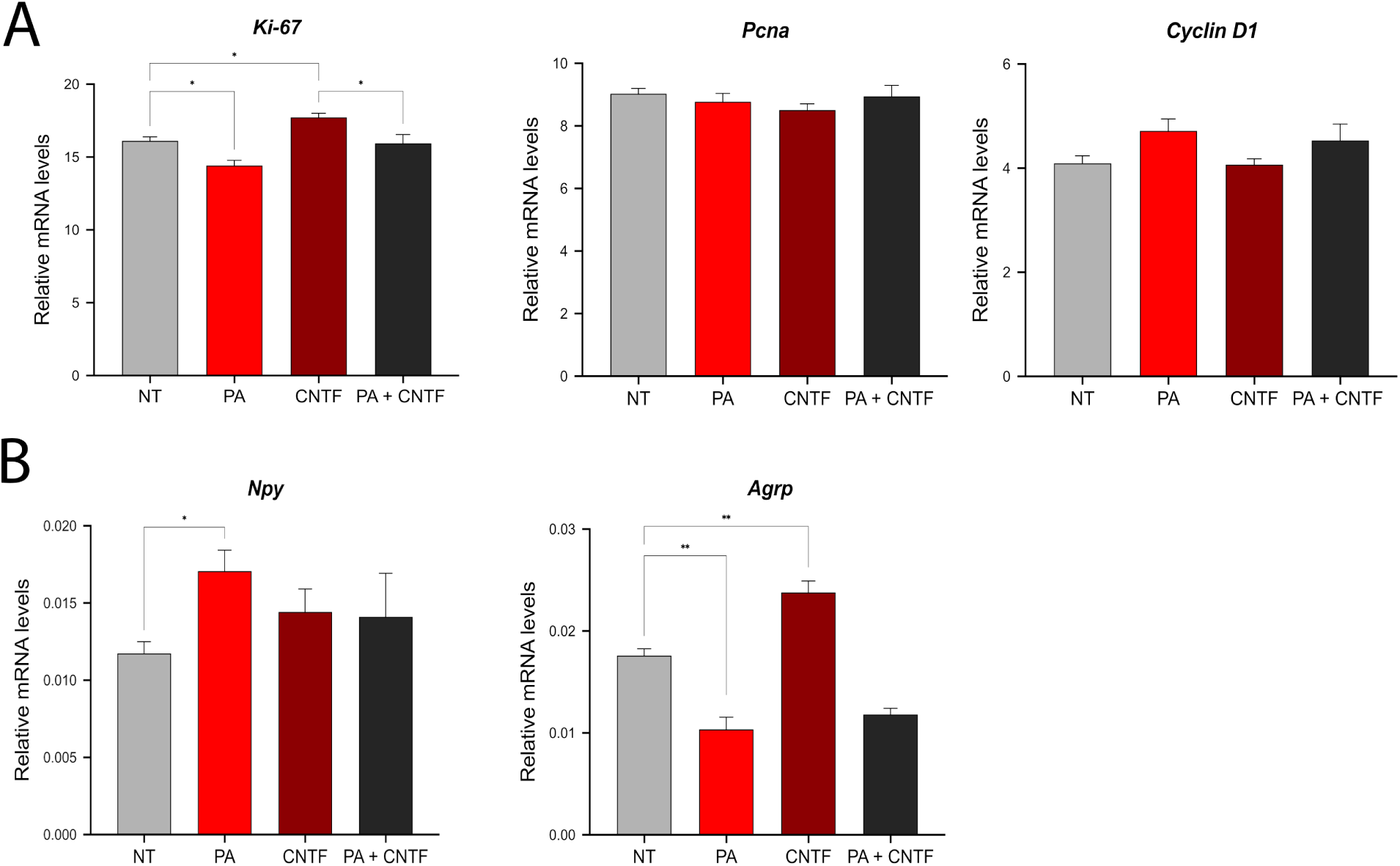
Proliferation-associated and orexigenic neuronal markers in N41 hypothalamic neurons under lipotoxic conditions. (A) Relative mRNA expression of *Ki-67*, *Pcna* and *Ccnd1* in N41 neurons treated with 200 μM PA, 1nM CNTF, or co-treated with PA and CNTF (PA + CNTF) for 24 h. (B) Relative mRNA expression of the orexigenic neuropeptides *Npy* and *Agrp* in N41 neurons treated with 200 μM PA, 1 nM CNTF, or co-treated with PA and CNTF (PA + CNTF) for 24 h. Gene expression was determined by RT-qPCR and expressed as relative mRNA levels (2^-ΔCt^) normalized to *Tbp*. Data are mean ± SEM from three independent experiments. *p < 0.05; **p < 0.01; ***p < 0.001, as determined by one-way ANOVA followed by Tukey’s post hoc test.

Because PA enhanced the expression of inflammatory mediators, including *Ikkβ*, and since activation of the IKKβ/NF-κB pathway has been associated with increased hypothalamic *Npy* expression^43^, we next investigated whether CNTF modulates the orexigenic neuropeptides *Npy* and *Agrp* under lipotoxic conditions. PA exposure for 24 h significantly increased *Npy* expression while reducing *Agrp* mRNA levels, highlighting an opposite regulation of the two neuropeptides in N41 neurons (Figure 7B). Combined CNTF and PA treatment did not modify the expression of either *Npy* or *Agrp* compared with PA alone (Figure 7B), indicating that CNTF does not counteract PA-induced changes in the expression of the two orexigenic neuropeptides.

## 4. Discussion

Beside its well-established peripheral metabolic effects, obesity is increasingly recognized as a condition in which chronic low-grade inflammation disrupts communication between peripheral metabolic organs and the brain, ultimately compromising the hypothalamic circuits involved in energy homeostasis^60^. Chronic overnutrition promotes ectopic lipid accumulation, leading to a persistent hypothalamic inflammation that impairs insulin and leptin signaling in key neuronal populations^32,61^. Reciprocal interactions between glial cells and neurons establish a feed-forward inflammatory loop that further exacerbates the hypothalamic dysfunction and contributes to systemic metabolic dysregulation^62^. Among the neuronal populations involved in metabolic regulation, NPY/AgRP neurons play a central role by integrating hormonal and nutrient-derived signals to coordinate feeding behavior, energy expenditure, and glucose homeostasis^63^. Consequently, obesity- associated hypothalamic inflammation alters AgRP neuronal function and responsiveness to peripheral metabolic signals, thereby contributing to whole-body metabolic dysregulation^32,61,63^.

Among the endogenous factors involved in hypothalamic energy homeostasis, CNTF has attracted considerable interest because of its potent anorexigenic and antidiabetic actions^19,37,64^. Experimental studies have shown that CNTF reduces food intake, promotes weight loss, improves glucose homeostasis, and enhances insulin sensitivity in both animal models and humans^19,37,64^. In the hypothalamus, CNTF activates canonical intracellular signaling pathways, including JAK2/STAT3 and MAPK/ERK, which are involved in energy homeostasis^35,36,38^. Besides such well-established metabolic actions, however, it is still largely unclear whether CNTF directly protects NPY/AgRP neurons against obesity-associated lipotoxic and inflammatory insults. Because neuronal responses in hypothalamic circuits are profoundly influenced by surrounding glial cells and neighboring neuronal populations, an *in vitro* model can provide insights into any direct actions of CNTF on NPY/AgRP neurons. We thus used N41 cells, a widely used *in vitro* model of NPY/AgRP-expressing hypothalamic neurons, to assess the effect of CNTF alone or combined with PA, a proven *in vitro* model of obesity-associated hypothalamic neuroinflammation and metabolic dysfunction. Our objective was to establish *i)* whether CNTF exerts any direct effects on hypothalamic neurons and *ii)* whether it is able to counteract PA-induced lipotoxicity.

Alone, CNTF activated the canonical intracellular signaling pathways JAK2/STAT3 and MAPK/ERK^2,47^, confirming the responsiveness of NPY/AgRP neurons to CNTF; it also upregulated the proliferation-associated markers Ki-67 and PCNA, consistent with its established neurotrophic properties, like promoting neuron survival *in vitro* and *in vivo*^4^ and maintaining adult neural stem cells^65^. The differential responses observed among Ki-67 and PCNA may reflect their distinct functions and temporal regulation during cell-cycle progression. Ki-67 is expressed throughout the active phases of the cell cycle, whereas PCNA is more closely associated with DNA replication^66^. Moreover, the increase in PCNA protein levels in the absence of detectable changes in *Pcna* mRNA may reflect differences in mRNA and protein kinetics, as well as post-transcriptional and post- translational regulatory mechanisms^67^. Thus, the lack of coordinated changes among these proliferation-associated markers likely reflects their distinct temporal and regulatory profiles following CNTF stimulation. Notably, previous studies have implicated CNTF in neuronal plasticity and adult hypothalamic neurogenesis^48,49^. This view is further supported by recent transcriptomic evidence that CNTF induces a pro-neurogenic transcriptional program in mouse subventricular zone- derived neurospheres cultures^50^. Although the biological significance of the upregulation of these proliferation-associated markers in our hypothalamic neuronal cell model remains to be established, our findings lend support to the neurotrophic role of CNTF and raise the possibility that it contributes to the maintenance and remodeling of hypothalamic neuronal networks. In N41 neurons, PA exposure induced a lipotoxic phenotype characterized by upregulation of inflammatory and ER stress markers, which are the key molecular features of obesity-associated hypothalamic dysfunction. CNTF significantly attenuated the PA-induced upregulation of key pro-inflammatory mediators, including *Mcp1* and *Tnfα*, supporting a protective role for CNTF against lipotoxic stress–induced neuroinflammation. Although the anti-inflammatory actions of CNTF in the central nervous system are not extensively characterized, CNTF does influence cytokine signaling and microglial responses in neural contexts by exerting immunomodulatory functions on neural tissues^68^. Our findings extend these observations to hypothalamic orexigenic neurons, indicating that CNTF can also blunt lipotoxic inflammatory responses at the neuronal level.

Given that inflammatory signaling is a major contributor to central insulin resistance, the anti- inflammatory effects of CNTF may contribute to preserve neuronal insulin responsiveness under lipotoxic conditions. In N41 neurons, PA exposure impaired insulin-stimulated AKT phosphorylation, consistent with the induction of insulin resistance under lipotoxic conditions, reported in hypothalamic neuronal models^41^. Notably, in PA-treated neurons CNTF restored insulin-induced AKT activation, suggesting that it preserves insulin signaling in states of metabolic stress. This observation is consistent with studies demonstrating anti-inflammatory actions of CNTF in peripheral metabolically relevant tissues, including human adipocytes, where CNTF reduced pro-inflammatory cytokine expression and improved insulin sensitivity^17^. Altogether, these findings suggest that CNTF not only curbs lipotoxic neuroinflammation but also preserves neuronal insulin responsiveness, supporting a role for it in hypothalamic metabolic homeostasis during obesity-associated metabolic stress.

Besides activating the JAK2/STAT3 and MAPK/ERK intracellular signaling pathways, CNTF also stimulated AMPK phosphorylation in NPY/AgRP neurons, extending previous observations made in peripheral metabolic tissues to hypothalamic neurons^17,19^. Concurrently, CNTF upregulated the genes associated with mitochondrial function and fatty acid metabolism, including *Ucp2*, *Cpt1α* and *Pgc1α*. Notably, *Ucp2* plays a key role in mitochondrial function regulation, reactive oxygen species production and substrate utilization. Indeed, *Ucp2* has been proposed as a metabolic switch regulating mitochondrial substrate selection^69^. In N41 neurons, CNTF blunted the PA-induced reduction of *Ucp2* expression while inducing a further increase in *Cpt1a* expression, suggesting that it preserves the metabolic adaptations that sustain mitochondrial function during lipid overload. Altogether, these findings indicate that, besides restraining inflammatory responses, CNTF may enhance the metabolic resilience of hypothalamic neurons by promoting mitochondrial adaptations that help maintain cellular energy homeostasis under lipotoxic conditions.

The present study has some limitations that should be considered when interpreting its findings. Although the N41 cell line is a valuable model for investigating the direct effects of CNTF on NPY/AgRP neurons, it clearly does not recreate the full complexity of the hypothalamic microenvironment. *In vivo*, NPY/AgRP neurons are embedded in highly interconnected neuronal circuits and maintain dynamic bidirectional communication with neighboring glial cells, particularly astrocytes and tanycytes, which actively participate in nutrient sensing, inflammatory responses and metabolic regulation^70^. Consequently, the isolated neuronal model used in this study cannot account for the contribution of neuron–glia interactions that shape hypothalamic responses to metabolic stress. Recent evidence has highlighted the central role of glial cells in orchestrating hypothalamic metabolic adaptations. In particular, elegant studies have demonstrated that microglial cells function as metabolic sensors and actively modulate hypothalamic neuronal activity, synaptic remodeling and whole-body energy homeostasis during high-fat feeding^70,71^. These observations suggest that the protective effects of CNTF identified in isolated NPY/AgRP neurons may be further shaped by neuron–glia interactions in the intact hypothalamic microenvironment. Studies employing more physiologically complex experimental systems, including neuron–glia co-cultures, hypothalamic organoids or *in vivo* models, are therefore required to determine how glial cells modulate the neuroprotective and metabolic actions of CNTF in the hypothalamic network. This notwithstanding, our study provides an essential first step toward dissecting the neuron-specific actions of CNTF.

Collectively, our findings identify CNTF as a direct regulator of NPY/AgRP neuron homeostasis. Beyond its established *in vivo* anorexigenic and metabolic actions, CNTF directly protects NPY/AgRP neurons from lipotoxic stress by simultaneously promoting pro-survival pathways, limiting inflammatory responses, preserving insulin signaling, and supporting metabolic adaptation. Our findings provide new insights into the cellular and metabolic actions of CNTF and further support its potential as a therapeutic agent in strategies aimed at counteracting obesity-associated hypothalamic dysfunction.

## Author contributions

**Jessica Perugini**: conceptualization, data analysis and interpretation, manuscript writing and financial support. **Edoardo Scopini:** methodology, performance of experiments, data analysis and interpretation. **Chiara Galli**: performance of experiments and figure preparation. **Martina Senzacqua:** performance of experiments and figure preparation. **Gaia Grandin**: performance of experiments. **Georgia Colleluori**: review and editing. **Ilenia Severi**: review and editing. **Antonio Giordano**: conceptualization, criticism during the study, review and editing and financial support.

## Acknowledgments

This work was supported by grants from Fondazione Cariverona (Bando Progetti Scientifici di Eccellenza 2018) to AG, EFSD Rising Star Grant 2024 to GC and Marche Polytechnic University (Contributi Ricerca Scientifica di Ateneo). The authors are grateful to Dr. Silvia Modena for her kind assistance with language and style.

## Data availability statement

The data that support the findings of this study are available from the corresponding author upon reasonable request.

